# Variability and Bias in Microbiome Metagenomic Sequencing: an Interlaboratory Study Comparing Experimental Protocols

**DOI:** 10.1101/2023.04.28.538741

**Authors:** Samuel P. Forry, Stephanie L. Servetas, Jason G. Kralj, Keng Soh, Michalis Hadjithomas, Raul Cano, Martha Carlin, Maria G de Amorim, Benjamin Auch, Matthew G Bakker, Thais F Bartelli, Juan P. Bustamante, Ignacio Cassol, Mauricio Chalita, Emmanuel Dias-Neto, Aaron Del Duca, Daryl M. Gohl, Jekaterina Kazantseva, Muyideen T. Haruna, Peter Menzel, Bruno S Moda, Lorieza Neuberger-Castillo, Diana N Nunes, Isha R. Patel, Rodrigo D. Peralta, Adrien Saliou, Rolf Schwarzer, Samantha Sevilla, Isabella K T M Takenaka, Jeremy R. Wang, Rob Knight, Dirk Gevers, Scott A. Jackson

## Abstract

**Background:** Several studies have documented the significant impact of methodological choices in microbiome analyses. The myriad of methodological options available complicate the replication of results and generally limit the comparability of findings between independent studies that use differing techniques and measurement pipelines. Here we describe the Mosaic Standards Challenge (MSC), an international interlaboratory study designed to assess the impact of methodological variables on the results. The MSC did not prescribe methods but rather asked participating labs to analyze 7 shared reference samples (5x human stool samples and 2x mock communities) using their standard laboratory methods. To capture the array of methodological variables, each participating lab completed a metadata reporting sheet that included 100 different questions regarding the details of their protocol. The goal of this study was to survey the methodological landscape for microbiome metagenomic sequencing (MGS) analyses and the impact of methodological decisions on metagenomic sequencing results.

**Results:** A total of 44 labs participated in the MSC by submitting results (16S or WGS) along with accompanying metadata; thirty 16S rRNA gene amplicon datasets and 14 WGS datasets were collected. The inclusion of two types of reference materials (human stool and mock communities) enabled analysis of both MGS measurement variability between different protocols using the biologically-relevant stool samples, and MGS bias with respect to ground truth values using the DNA mixtures. Owing to the compositional nature of MGS measurements, analyses were conducted on the ratio of Firmicutes: Bacteroidetes allowing us to directly apply common statistical methods. The resulting analysis demonstrated that protocol choices have significant effects, including both bias of the MGS measurement associated with a particular methodological choices, as well as effects on measurement robustness as observed through the spread of results between labs making similar methodological choices. In the analysis of the DNA mock communities, MGS measurement bias was observed even when there was general consensus among the participating laboratories.

**Conclusion:** This study was the result of a collaborative effort that included academic, commercial, and government labs. In addition to highlighting the impact of different methodological decisions on MGS result comparability, this work also provides insights for consideration in future microbiome measurement study design.

## Introduction

Over the last decade, advances in DNA sequencing technology (Next-Generation Sequencing or NGS) have led to its widespread adoption by the scientific community for myriad applications. One such application, known as metagenomic sequencing (MGS), has led to a transformation in how we measure and characterize complex microbial communities of microbiomes. MGS has emerged as an important and powerful tool as we seek to comprehend the roles of microbes inside complex and dynamic communities that are both capable of maintaining and harming human and environmental health. MGS measurements are able to ‘see’ whole classes of microorganisms present in a microbiome sample (e.g., all bacteria by 16S rRNA gene amplicon sequencing (16S), or all dsDNA by whole-genome shotgun WGS); MGS can also assign a relative abundance to each microorganism in complex samples. [1-4] Because of these advantages, MGS is being increasingly adopted across diverse application spaces including infectious disease diagnostics, [5-12] epidemiological investigations, [13-15] food safety, [16] and biothreat surveillance. [9, 17-19]. The results of MGS measurements have been used to diagnose infectious diseases that were missed by conventional methods.[20, 21] As such, regulatory agencies are actively developing new guidance and policies regarding the use of MGS in the clinic and in other regulated spaces.

While MGS measurements hold great promise in monitoring and understanding microbial communities, the current impact is often hampered by a lack of reproducibility and comparability, particularly between different research centers. [22-24] MGS measurement results are the product of complex workflows incorporating multiple distinct steps and involving a multitude of methodological choices (e.g., sample collection and storage, DNA extraction and purification, NGS library preparation either for WGS or 16S, DNA sequencing platform, data cleanup and processing, bioinformatic analysis, interpretation). Throughout this workflow, measurement bias (deviation from ground truth) and measurement noise (experimental variability) are potentially introduced with each step and will depend on the particular methodological choices made.[25] It is widely recognized that the interlaboratory reproducibility of MGS microbiome measurements is poor, and there have been numerous efforts aimed at benchmarking the analytical performance of MGS measurements in terms of sensitivity, specificity, precision, reproducibility, etc. [26-33] These challenges are well-documented, and the community has long recognized the need for studies to prioritize and investigate the sources of variability and bias in the experimental workflow [27] and the need for standardized materials and methods to improve the comparability and scope of MGS measurement results.

Designing the studies to identify sources of variability and bias as outlined above comes with its own set of challenges including: sufficient numbers and diversity of reference samples to help power the study; testing of a wide range of variables; a lack of consistent data analysis; cost & coordination. While the task may seem daunting, several groups have taken up the call to begin to address these challenges. In recognition of the complexity of the workflow, some groups have broken the MGS workflow into more manageable sections with most of the focus being directed at characterizing the effect of data processing and analysis either using *in silico* datasets [32, 33] or metagenomic DNA control material [26, 28, 34-38]. Other groups have sought to capture bias throughout the workflow by distributing sets of identical microbiome samples [39, 40].

Herein, we describe the Mosaic Standards Challenge (MSC). The MSC brought together academic, federal, and private industry partners in an international interlaboratory study focused on capturing the diversity of protocols and methodological choices involved in NGS-based microbiome measurements and understanding their impact on observed taxonomic profiles. To achieve this, we produced a panel of homogeneous microbiome samples, developed a custom cloud-based web portal for collecting sequencing data and metadata, [41, 42] and statistically evaluated the MGS results. The microbiome samples included human feces from multiple donors and DNA mock communities. For every sample analyzed, nearly 100 metadata parameters describing the MGS protocol were collected, with participation from 44 MGS laboratories. The resulting analysis demonstrated that various protocol choices have significant effects that range from skewing MGS measurement results (e.g., WGS or 16S analysis) to increasing measurement robustness (e.g., homogenizer use during DNA extraction). The ground truth DNA mock community samples revealed that MGS measurement bias can persist, even when there is consensus (measurement agreement) among results from different laboratories.

## Results

Briefly, the study consisted of three components: reference material selection and production, broad participation from the microbiome community including metadata reporting and MGS data uploads, and common analysis pipelines applied to the raw sequencing data alongside the methodological metadata from each participating laboratory. The timeline and overall workflow of the MSC are shown in Figure 1.

**Figure 1.**
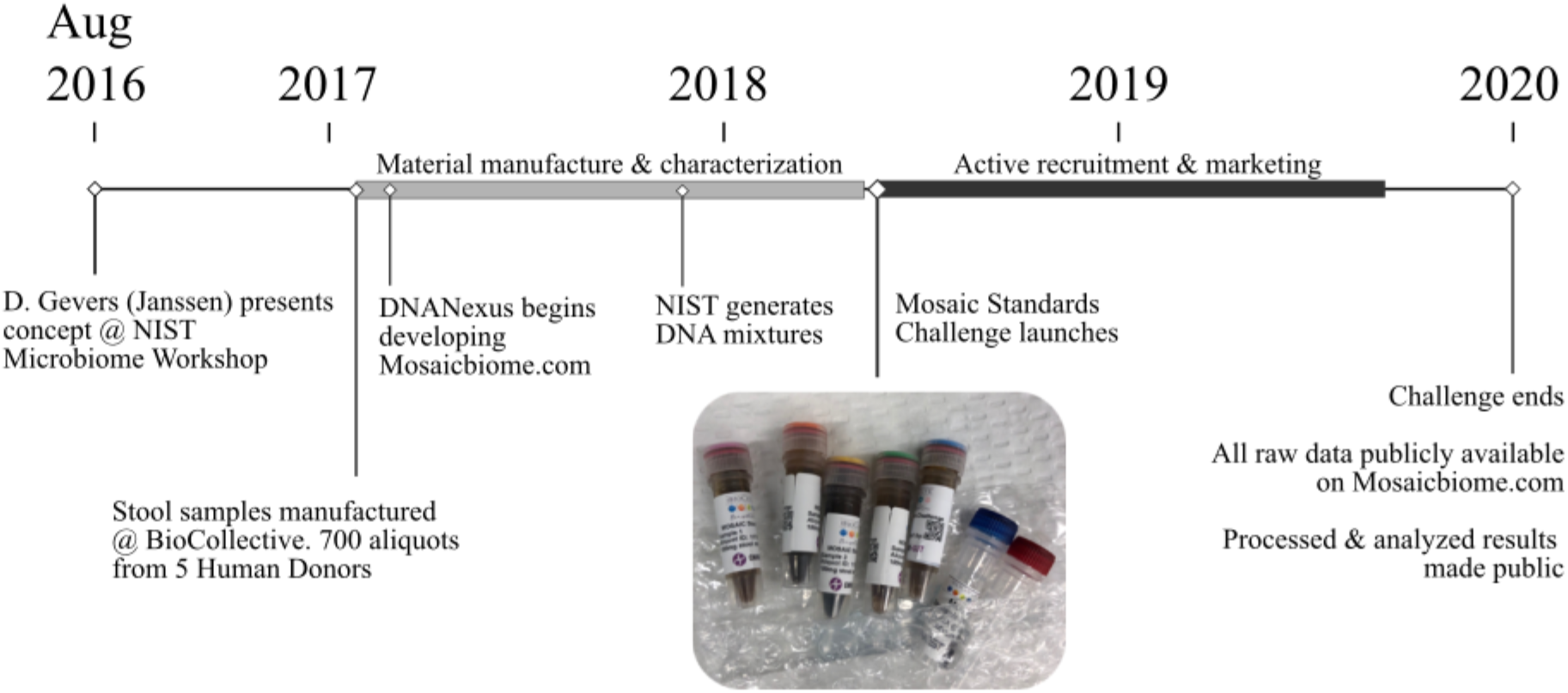
Study design timeline. Inset image shows material received by participants.

### Material Production

The reference samples selected and distributed in this study included 5 human stool samples and 2 DNA mixtures (mock DNA communities). The five stool samples were selected from a pool of potential donors based on the dissimilarity of their microbiome composition (Figure 2).

**Figure 2.**
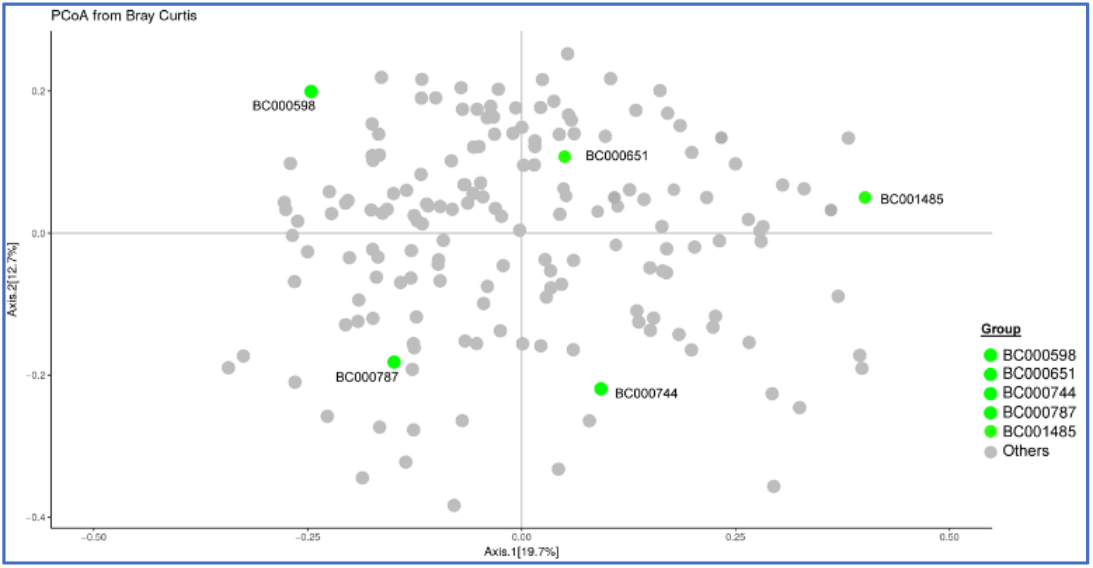
Principal coordinate analysis of donor samples in the BioCollective stool collection. Stool samples included in the Mosaic study (green points) were selected based on their PCoA diversity within the constellation of samples available from the BioCollective. All selected donors self-reported being healthy except BC001485, who reported Parkinson’s Disease.

For each sample, multiple stool donations from an anonymous individual were homogenized in the presence of a stabilization buffer to produce 1-liter of homogenized, stabilized fecal material. Two allochthonous microorganisms, *Aliivibrio fischeri* and *Leifsonia xyli*, were also added to each batch of stool (∼10^8^ cells/mL) and homogenized. Approximately 700 aliquots (1 ml per aliquot) were prepared from each of the 5 batches, and aliquots were stored at -20 °C until ready to ship to participants. To verify that these materials were sufficiently homogenous, 10 aliquots were selected randomly from each of the 5 batches and subjected to both 16S and WGS analysis. These results (Figures 3 and Si-1) indicate that (i) each individual sample donor has a unique microbiome composition, and (ii) the stool samples are suitably homogenized (fit-for-purpose).

**Figure 3.**
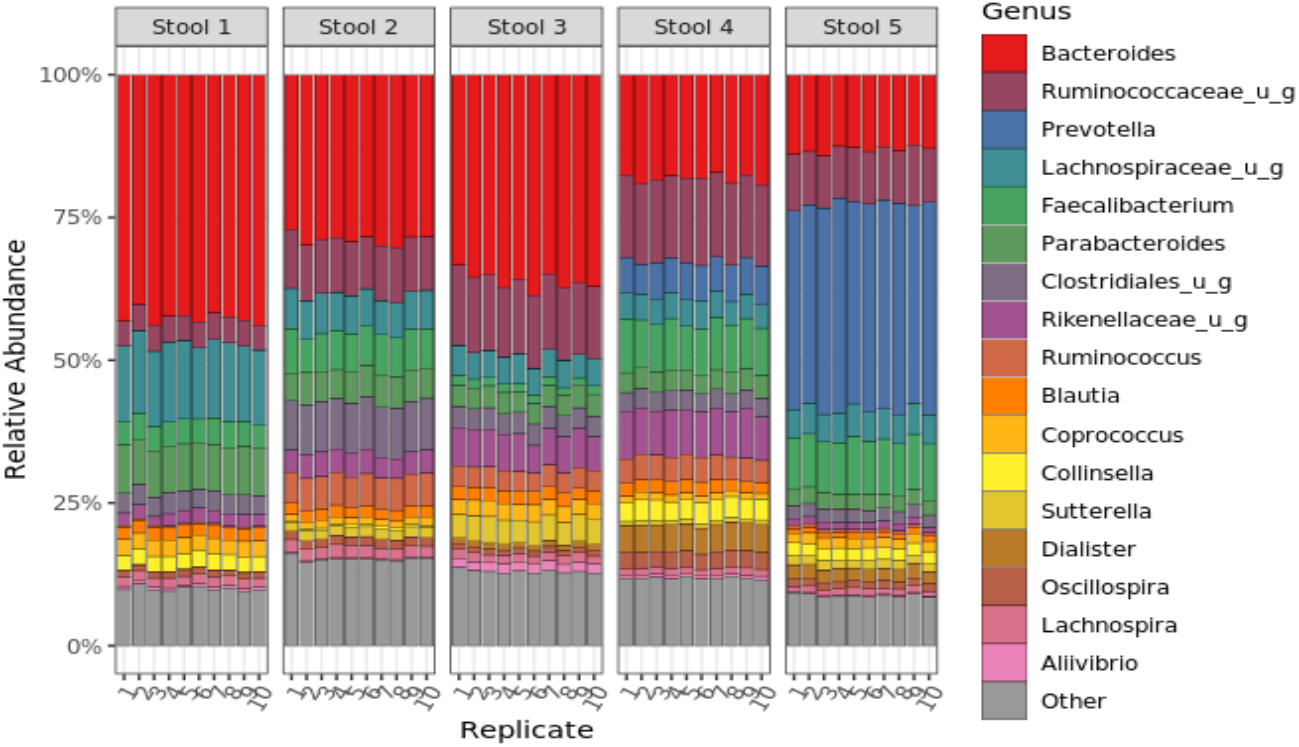
Metagenomic sequencing analysis of Mosaic stool samples to determine homogeneity of samples. The bar chart shows the relative abundance as measured by 16S rRNA MGS at the genus level for 10 replicate tubes from each stool sample (stool 1-5). Taxa colors denote the 17 most abundant genera overall, as well as an exogenously added internal standard; all other genera are grouped as ‘other’ and shown in grey. MGS analysis by WGS also exhibited good homogeneity (Figure SI-1).

Additionally, 2 DNA-based mixtures were prepared for the MSC where ground-truth taxa abundances could be assigned. Both materials (Mix A and Mix B) were mixtures of genomic DNA (mock communities) that were extracted from pure cultures of 13 bacterial species mixed at roughly equal genomic ratios (Mix A) or at varying abundances across 3-orders of magnitude (Mix B).

### Recruitment and Community Participation

To kick-off the study, a targeted media campaign was launched to recruit participation in the MSC; study enrollment was open from May 2018 until December 2019. [43, 44] Each lab that volunteered to participate was shipped the 5 stool samples and two DNA samples free of charge. By design, the MSC did not prescribe any required methods or instrumentation to the participants. Rather, participants were instructed to use their own in-house protocols and encouraged to explore new methods. To capture these methodological details, a comprehensive standardized metadata reporting sheet was developed and deployed where participants could record the details of their protocols. This metadata reporting form included over 100 questions and was intended to capture the most intricate details of each step in the measurement process (The metadata capture questions are available in the supplemental file 1.) Both methodological data and raw data were then captured using a custom web-based cloud analytics portal that enabled the collection, storage, analysis, and visualization of MGS data generated by the MSC participants [41]. This not only facilitated analysis within a single bioinformatics pipeline; it also enabled participants to view their results in the context of all other MSC results immediately following upload. Thus, participants could quickly visualize how their methods compared to others in the community.

A total of 44 labs participated in the MSC by submitting MGS results (16S or WGS) along with accompanying metadata (Table 1). Most labs analyzed all samples, though some only analyzed the stool samples. Of the 44 MGS submissions, 30 were 16S rRNA datasets and 14 were WGS datasets (Table 1). On average, 16S rRNA MGS datasets had ≥10^5^ reads, while the WGS analyses were typically a log higher with >10^6^ reads (Figure SI-2). Significant variation in read number was observed both between participating labs and individual samples (Figure SI-2).

**Table 1.** A total of 44 labs submitted 16S or WGS analyses of the Mosaic Stool and DNA samples with accompanying metadata. Two sequencing datasets with incomplete metadata were dropped.

| <u>MGS Analysis</u> | <u>Samples analyzed</u> | <u>Number of Labs</u> |
| --- | --- | --- |
| 16S | 5x Stool | 8 |
|  | 2x DNA | 0 |
|  | 7x (Stool and DNA) | 22 |
| WGS | 5x Stool | 0 |
|  | 2x DNA | 0 |
|  | 7x (Stool and DNA) | 14 |

### Metagenomic Sequencing (MGS) Interlaboratory Comparison

A Bray-Curtis principal coordinate analysis (PCoA) for both the 16S (n=150) and WGS (n=70) datasets (Figure 4a and 4b) demonstrate that the biological variability (i.e., stool sample ID) was the major factor influencing the overall ordination of the data, as expected. The impact of methodological variability can be seen via the dispersal of datasets within each stool sample. It’s noteworthy that the outputs from the 16S datasets and the WGS datasets were so dissimilar that separate PCoA analysis plots were required. From the PCoA plot of the 16S data (Figure 4a), we observed that one of the participating labs made an apparent transposition in the labeling of samples 3, 4, and 5. Based on this apparent error, we excluded all the data (stool samples 1-5) from this lab for the remainder of the analyses described in this manuscript.

**Figure 4:**
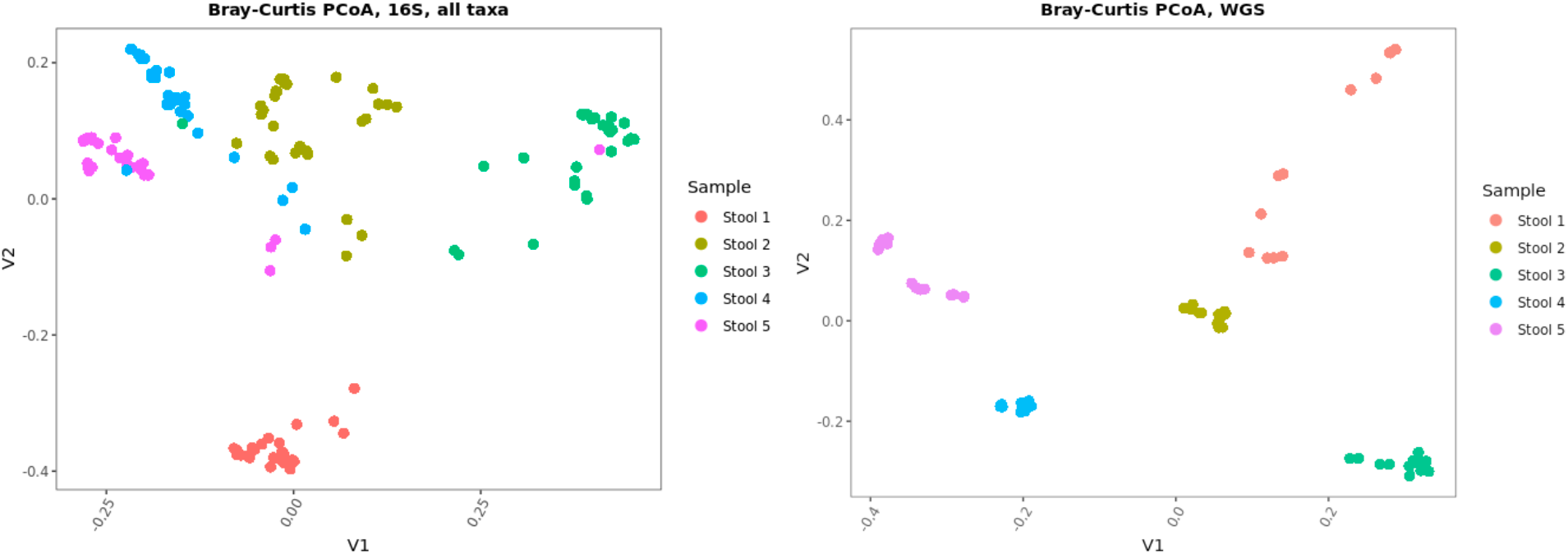
Principal coordinate plots of the Bray-Curtis dissimilarities for 16S and WGS analyses exhibits clustering by Stool sample. Each data point represents a distinct laboratory analysis of each sample. The separation in the clusters is attributed to methodological differences between labs.

#### Firmicutes:Bacteroidetes ratio

Because of the compositional nature of MGS results, individual taxa relative abundances are not directly comparable between different samples.[45, 46] Instead, ratios of taxa within each sample were expected to be more reliable because the effects of sample composition on each taxa relative abundance could cancel out. One ratio that has been of interest in the field is the ratio of phyla Firmicutes:Bacteroidetes; therefore, we chose this ratio to demonstrate the utility of using ratios of taxa to compare data between samples [39-42]. Thus, this ratio was utilized and included in our results purely for its bioinformatic utility and is not intended to serve as an indicator of gut health or dysbiosis. The Firmicutes:Bacteroidetes ratio was calculated for each Mosaic stool sample and compared among the individual laboratory results (Figure 5). As was expected since each laboratory used their individual MGS protocols (e.g., methodological choices for DNA extraction, library preparation, and sequencing), the Firmicutes:Bacteroidetes ratio varied substantially both between stool samples within each lab, as well as between labs.

**Figure 5.**
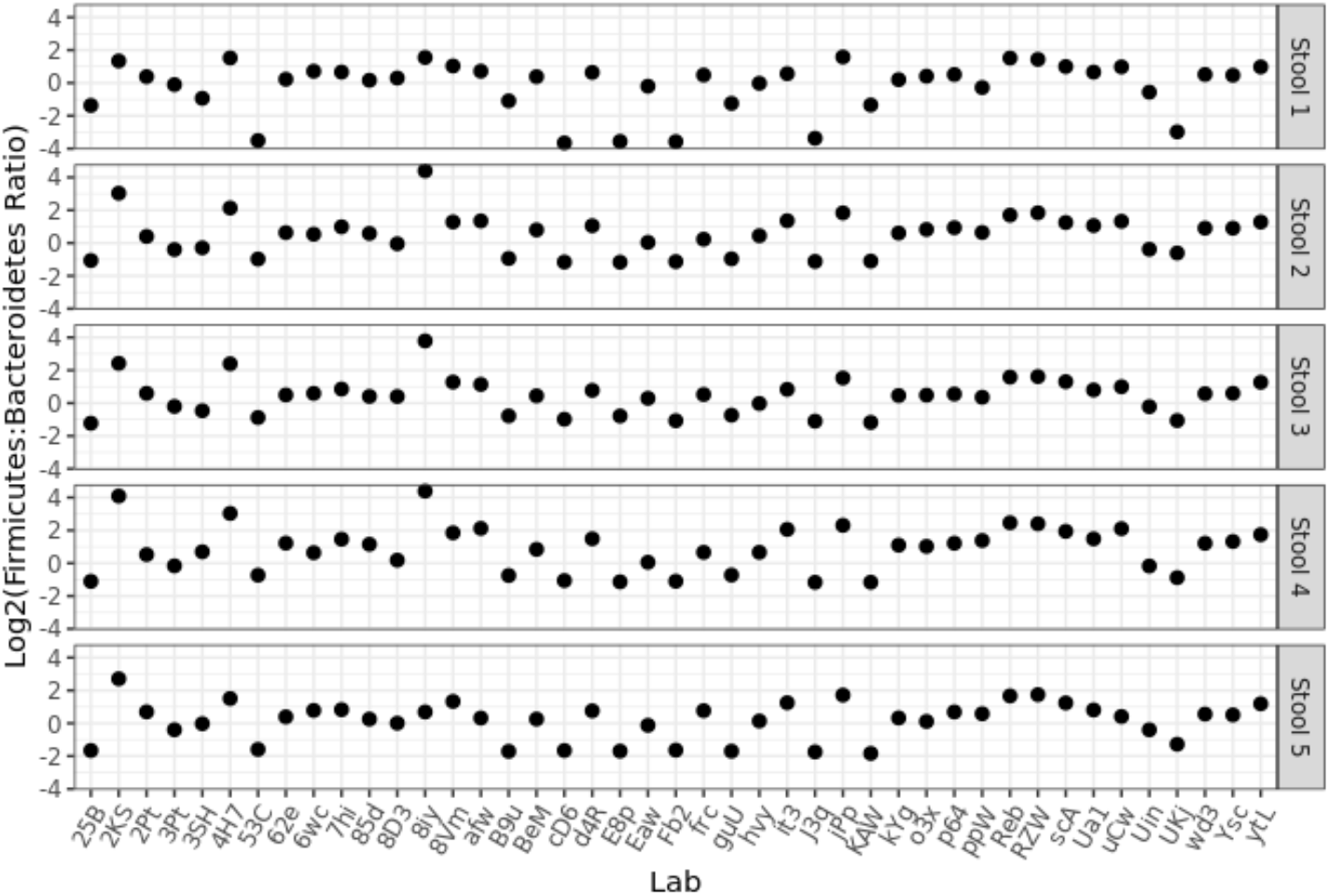
The Firmicutes:Bacteroidetes ratio was calculated for all stool samples and plotted for each participating laboratory. Of note, data submission was anonymous, so multiple submissions from the same research center would appear as distinct labs.

#### Amplicon vs. Shotgun sequencing

One goal of the MSC was to determine how the selection of different methodological parameters during MGS would lead to observed differences in the taxonomic profiles and relative abundances. The highest-level methodological choice was between 16S MGS or WGS MGS. Indeed, the Firmicutes:Bacteroidetes ratio was affected by the type of sequencing performed, with 16S MGS analyses reporting significantly higher Firmicutes:Bacteroidetes ratios (Figure 6a). While the majority of the 16S MGS datasets indicated that Firmicutes were present at a higher relative abundance than Bacteroidetes, WGS data found the inverse with Bacteroidetes being present at a higher relative abundance than Firmicutes. The magnitude of this effect was quantified by averaging the results from all labs reporting each methodological parameter (e.g., 16S or WGS for sequencing strategy) divided by the average result overall and plotted as a fold change on a log scale (Figure 6b). The dependence of the Firmicutes:Bacteroidetes ratio on analysis strategy that was observed in this dataset could explain recent reports that question the reliability of the Firmicutes:Bacteroidetes ratio as a diagnostic indicator of gut health [47]. This dependence was consistent across all five stool samples (Figure SI-3), and an analysis of each sample’s alpha diversity also yielded a similar stratification with respect to the methodological choice of 16S or WGS analysis (Figure SI-4).

**Figure 6.**
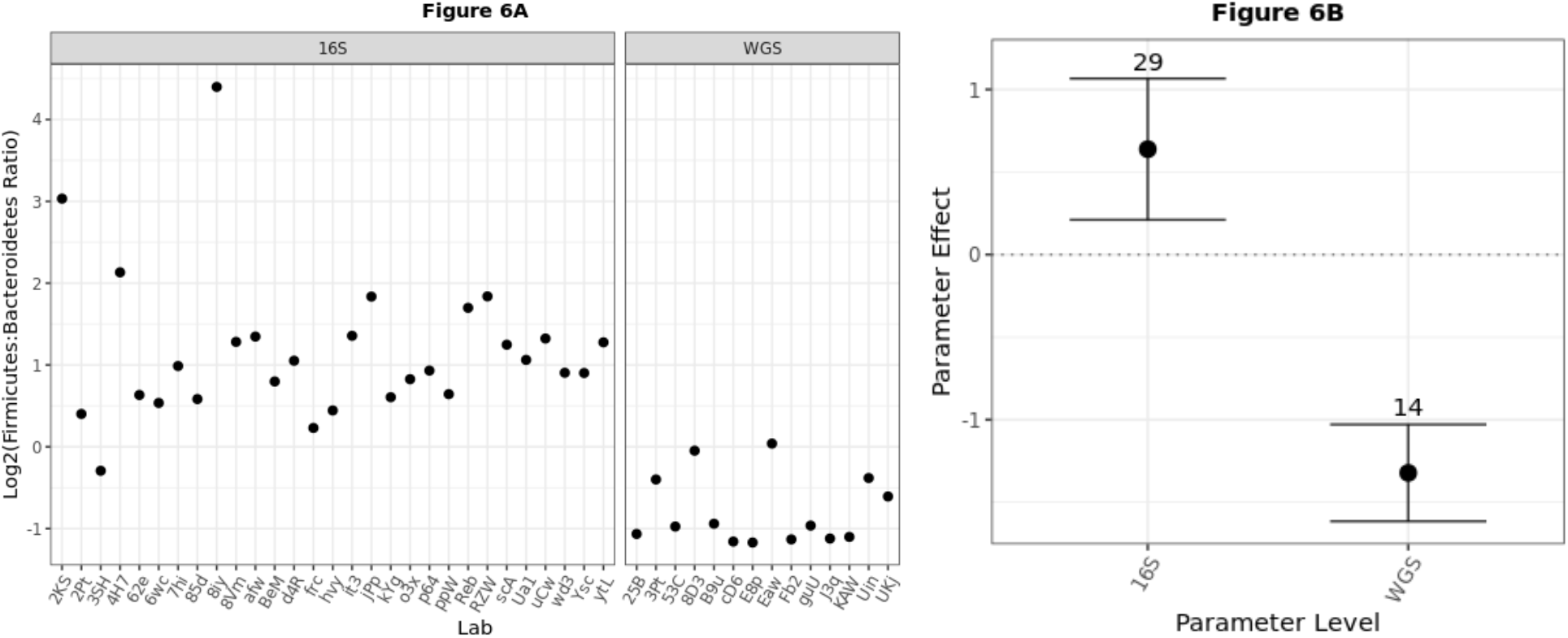
The effect of analysis strategy (16S versus WGS) on the Firmicutes:Bacteroidetes ratio was readily observed for just one stool sample by simple grouping (A), and the effect was quantified (B) by dividing the average results among labs reporting the specified parameter level by the average results overall. In B, this parameter effect was plotted on a log (base 2) scale, such that the horizontal line at 0 denotes the null hypothesis of no effect; error bars show the 99% confidence interval. Quantified effects for the other stool samples were similar and are included in Figure SI-3. Similar stratification was observed when measuring each sample’s Inverse Simpson alpha diversity (Figure SI-4).

#### Other metadata parameters

When submitting results, participating labs were asked to complete a standardized metadata reporting sheet that included 100 different questions regarding the details of their protocol. Some questions were generally applicable like “what sequencing instrument did you use” while others were more nuanced like “what was the PCR primer set used.” As such, some fields were required, and others were optional. Because of the large impact generated by the 16S vs. WGS methodological variable (Figure 6) and the hierarchical nature of other methodological choices (e.g., ‘What was the target gene amplicon’), we chose to analyze each data set separately. The effect on the Firmicutes:Bacteroidetes ratio on the 16S MGS results was quantified for each subsequent methodological choice (Figure 7) in a similar manner to that employed in Figure 6b. While there were many methodological variables that appeared to have a significant impact on the results (Figure 7; similar analysis for other stool samples is included in Figure SI-5), many of these were only reported by a single lab (n=1). Of the 30 labs submitting 16S MGS data, there were 14 methodological differences in their protocols. Of the 14 labs submitting WGS data, there were 9 methodological differences in their protocols (Figure SI-6). Not all methodologic variables had a significant impact on the result. Methodological variables that were observed to have a significant impact on the 16S MGS results for 2 or more stool samples (parameter effect and 99% confidence interval) included the manufacturer of the DNA extraction kit and the target gene for amplification (Figure 7 and SI-5). Methodological variables that were observed to have a significant impact on the WGS results (parameter effect) for 2 or more stool samples included the DNA extraction protocol, the manufacturer of the DNA extraction kit, and the library kit for shotgun sequencing. In addition to their impact on the parameter effect as described above, some methodological variables were observed to have a significant impact on the robustness of the measurement (observed as a lack of variability when other parameters are varied). For example, when asked whether a homogenizer or shaking apparatus was used, those labs that reported “Yes” displayed much greater robustness than those reporting “No” (Figure 7 and SI-5 and SI-6).

**Figure 7.**
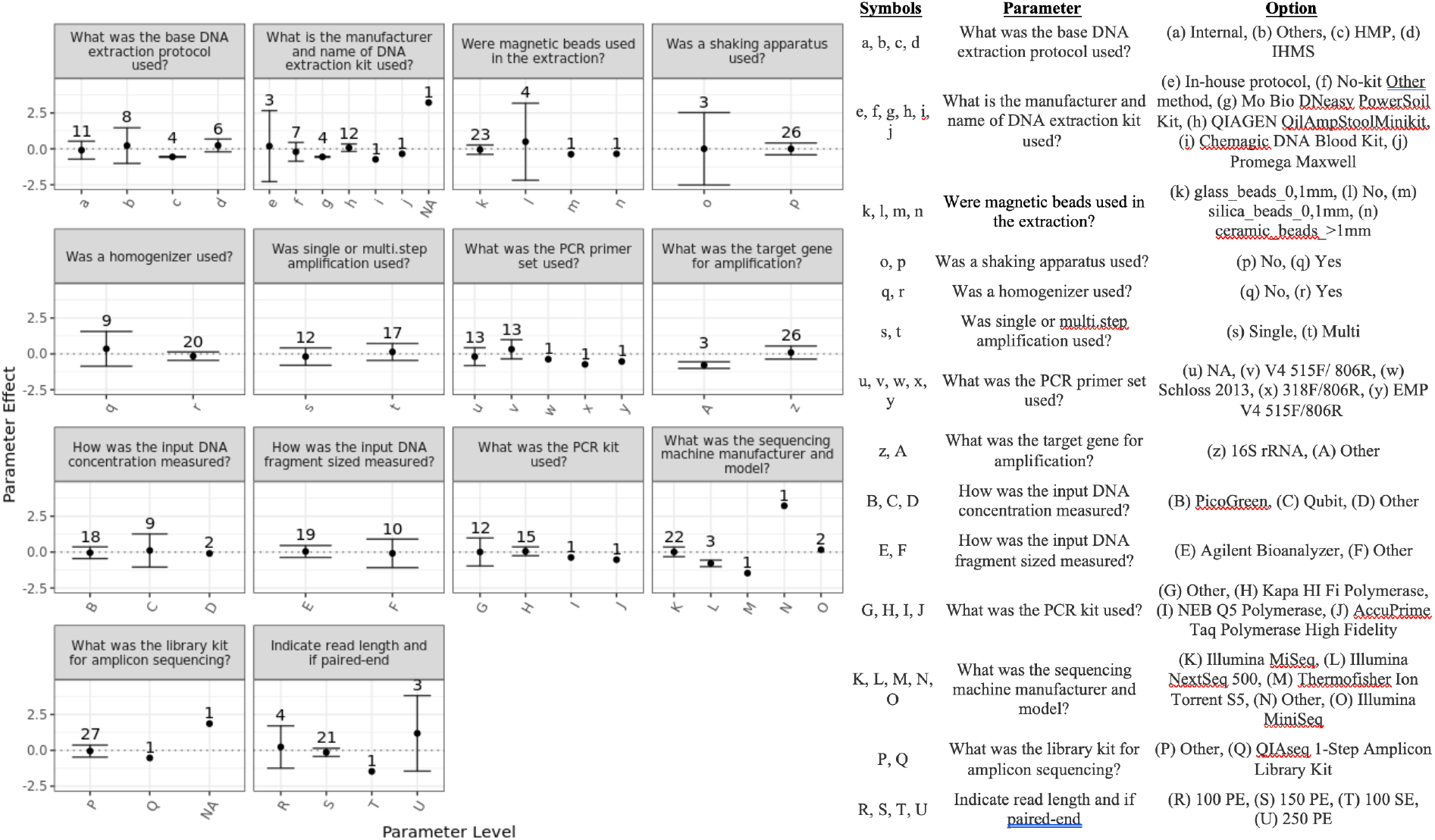
Within labs performing 16S amplicon sequencing, the parameter effect on the Firmicutes:Bacteroidetes ratio was calculated as described in Figure 6 for each relevant metadata parameter. Shown here from just one stool sample, results from other stool samples were similar and are provided in Figure SI-5.

#### ‘Spike-in’ organisms

An additional attribute of the fecal materials used for this interlaboratory study was the inclusion of two exogenous organisms to serve as whole-cell internal controls (i.e., spike-ins). Since these organisms were added during the bulk homogenization step, their abundance should be constant across all the stool sample aliquots. As such, it was expected that the ratio of *A. fisherii* to *L. xyli* would be constant for each particular methodology (e.g., within a lab).

Surprisingly, *L. xyli* was not identified in any of the submitted 16S datasets and was only observed at a low abundance (approximately 0.001 %) by WGS analysis. When the *A. fisherii*:*L. xyli* ratio (by WGS) was plotted for each participating laboratory (Figure SI-7), significant variability between samples was observed. These data were unexpected and could have resulted from poor database representation of *L. xyli* in the commercially available bioinformatic pipeline used, inefficient DNA extraction, or low or inconsistent distribution during material manufacture, among other possible explanations.

#### Genomic DNA mixtures

Another control included in the interlaboratory study were mixtures of purified microbial genomic DNA. These were included alongside the stool samples in the Mosaic Kit to serve as parallel processing controls and included two different mixtures, one equigenomic between taxa (Mix A) and one with ten-fold dilutions of the various taxa (Mix B). These genomic DNA mixtures were validated for genome copy number using ddPCR (droplet digital PCR) and serve as ‘ground truth’ for the MGS measurements. For comparison to the MGS measurements, genome copy number (as measured by ddPCR) was scaled by the assembled genomes of the individual strains (i.e., rRNA copy number or genome size) to yield ground truth values for comparison to 16S or WGS results, respectively. As with the Firmicutes:Bacteroidetes ratio described above, ratios of individual taxa were used to characterize the DNA mock communities and remove the compositional dependence of the raw relative abundance assignments. Since these analyses included 16S sequencing results, we focused on strains that were unique at the genus level, yielding 6 distinct ratios within each sample. The independent determination of actual DNA concentration (ddPCR) was compared to the results of MGS analyses (Figure 8).While there was some agreement among participating laboratories (consensus), their results generally differed from the actual abundances. Overall, this indicates that even when consensus exists among MGS results, significant unidentified bias can remain. Further, this was taxa-dependent, with some taxa (e.g., Kp by 16S or Pa by WGS) producing particularly significant variability and deviation from ground truth.

**Figure 8.**
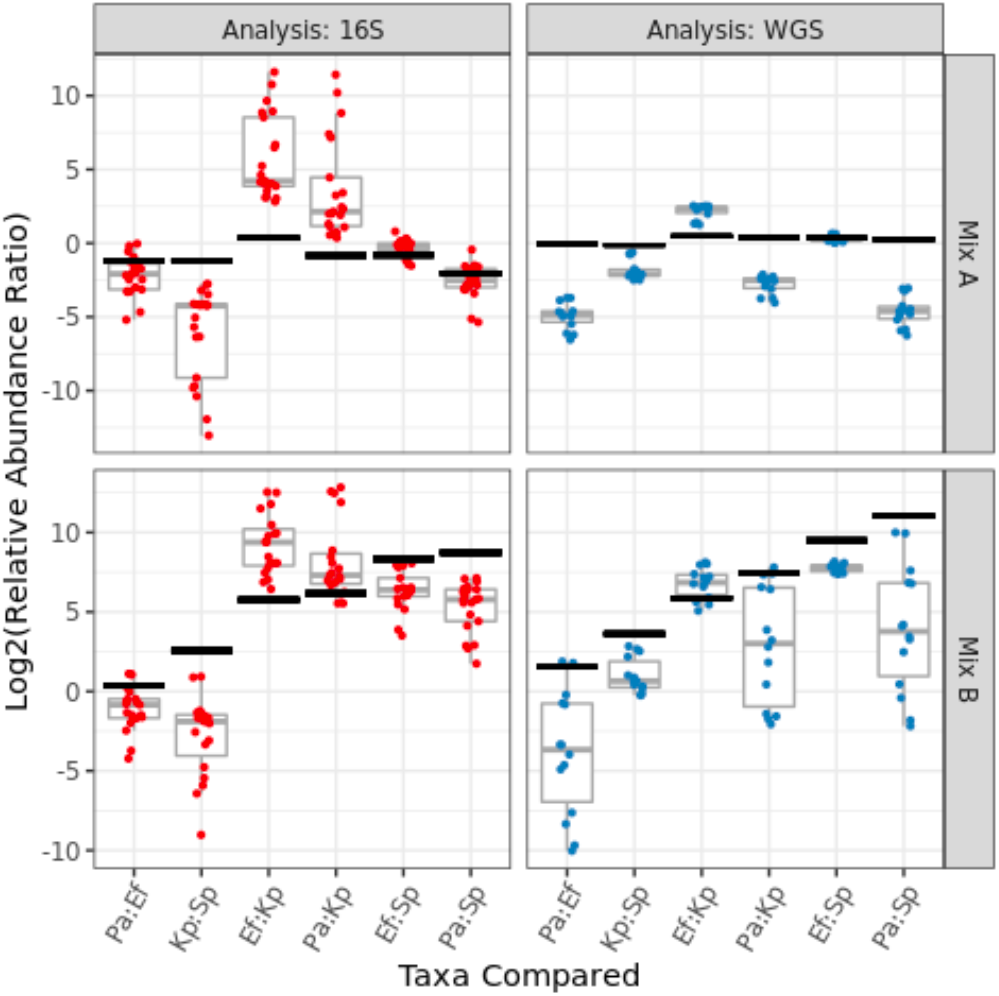
For the DNA mixtures, independently measured ‘ground truth’ results (black 99% confidence intervals) for the ratios between taxa relative abundances can be compared to each individual lab’s amplicon (red points) or shotgun (blue points) metagenomic sequencing results, as well as the range of results (grey boxplots) among participating labs. The taxa in Mix A were roughly equally abundant, while the Mix B sample exhibited groups of taxa added at 10-fold dilutions. The horizontal axis identifies the taxa (known to be present in the DNA mixtures) whose observed relative abundances were ratioed. The ground truth values were scaled to account for known 16S copy numbers (for amplicon sequencing) or genome sizes (for shotgun sequencing), so the ‘actual’ ratios vary slightly between the two analyses even though the DNA concentrations are identical. Genus-level taxonomic bar charts by (16S and WGS analysis) show the average composition observed for each DNA mixture (Figure SI-8).

## Discussion

The MSC represented the third in a series of community challenges of increasing complexity hosted by Janssen’s Human Microbiome Institute (JHMI) as an effort to improve the overall quality of microbiome MGS measurements. This study was designed and implemented through a collaborative effort that included the Janssen Human Microbiome Institute (JHMI), The BioCollective, LLC (TBC), DNAGenotek, DNANexus, and the National Institute of Standards and Technology (NIST) which serves as the National Metrology Institute for the U.S. These organizations in turn represent biopharmaceutical companies, biotechnology companies, data analytics companies, and Federal Government laboratories, all of whom have a vested interest in reliable and comparable microbiome measurements. The goal of the MSC was to capture the diversity of protocols for MGS-based microbiome measurements in an effort to begin to elucidate the impact of these methodological variables on the resulting taxonomic profiles and guide the development of future reference materials.

The MGS workflow required for microbiome analyses is complex. Therefore, designing an interlaboratory study that includes a multitude of the methodological variables and assesses their effect on the results is an ambitious project. Several teams have sought to address the question of methodological bias and variability over the years. [26-28, 31-33, 39, 40] These investigations have taken a variety of approaches from prescribing locked-down SOPs and analyzing specific samples to more open-ended data collection. The interlaboratory study presented herein specified seven samples for analysis (5 different stool samples and 2 DNA mixtures) while intentionally leaving protocol choices up the participating labs, both to sample a diverse set of methodological parameters as well as to survey common methodological choices.

The design and implementation of this project can be broken into three major areas: (1) reference material selection and production, (2) capturing metadata and MGS raw data, and (3) comparing results between participating laboratories.

### Reference Material Production

One of the first decisions was the identification of reference material(s) to include. There are two primary types of materials that have been used for this type of study: (1) biologically derived microbiome samples and (2) mock communities. Both types of materials were included in the current investigation because they are useful in different ways for comparing between diverse analytical workflows.

For biologically derived microbiome reference materials, a natural community (e.g. sludge, soil, fecal material) is collected, homogenized, and aliquoted. Previous interlaboratory studies have used these homogenized real-world materials [31, 39]; however, the number of units needed and the associated costs of a large-scale study are often prohibitive. Further, while biologically-derived materials represent the complexity and diversity of real-world samples, they currently lack ground-truth value assignments (e.g. actual taxonomic abundances) due to a lack of unbiased analytical methods (e.g., DNA extraction, PCR amplification) and the inherent ambiguity associated with microbial taxonomy that hinders our ability to define clear measurands (e.g., Escherichia coli vs. Shigella, or the recent reclassification of Lactobacillus into 23 novel genera). [48] The addition of allochthonous bacteria (“spike-ins”) at consistent abundances into biologically-derived materials can provide some ground truth values to facilitate the assessment of MGS measurements.

Nevertheless, these biologically-derived materials remain useful for comparing methods and assessing measurement precision within individual laboratories and across different laboratories. In the current study, five stool samples were selected based on their dissimilarity from one another among a constellation of potential stool donors (Figure 2), with the intention of representing the variability of naturally-occurring samples. Preliminary in-house analysis of individual aliquots demonstrated (Figure 3) that the material collection and preparation resulted in samples with reliable between-aliquot homogeneity, even given the inherently inhomogeneous starting point of multiple donations of human stool.

Mock community reference materials are laboratory-prepared mixtures of defined constituents (typically DNA from individually cultured bacteria; sometimes mixtures of whole cells) at specified amounts. Thus, these materials are useful as ‘ground truth’ for analysis workflows, allowing quantitative assessment of analytical performance (e.g., accuracy, bias, precision, etc.). However, these mock community materials are inherently non-biomimetic of actual microbiome samples (e.g., feces, soil, etc), namely due to their low complexity and the absence of a matrix-effect, which can limit their utility for assessing analysis workflows. [26] By including both types of reference materials (5x stool samples and 2x mock communities) in the MSC, we sought to include the benefits of both, using the biologically-derived materials to assess MGS measurement variability between different protocols, and using the DNA mixtures to assess MGS bias with respect to ground truth values.

### Capturing Metadata

The universe of discrete MGS methodologies is quite large. Preliminary projections during project planning estimated that several hundred samples would be needed to fully explore this methodological space. Thus, the MSC set-out to host an international interlaboratory study on an unprecedented scale. 700 units of reference material were prepared and made available free-of-charge, where each unit consisted of 5 distinct, biologically-derived human fecal microbiome samples and 2 DNA mixtures (mock communities). To our knowledge, there has never been an MGS interlab study designed on such a massive scale. However, during the 19-month marketing campaign, only ∼100 units were distributed. Further, from these recipients, only 44 sets of raw sequencing data and metadata were submitted (Table 1), limiting the statistical power of the resulting analyses. Nevertheless, the unused units remain currently available from The BioCollective, allowing interested researchers to analyze with their own methods using the same samples that have been characterized and reported on here.

Alongside the raw sequencing data submitted, participating laboratories filled out a metadata questionnaire (available in supplemental file 1) with ∼100 discrete questions about the methods employed, most of which allowed selection from drop-down options describing the most common methodological choices. However, it must be noted that even these in-depth options were not sufficient to encompass all experimental possibilities, and many metadata selections represented ‘Other’ or ‘Internal Method’ options. And, of course, the number of potential methodologies continues to expand as new techniques are developed or made commercially available. It was also apparent within the submitted metadata that the observed methodological choices were not randomly distributed. There was no effort made in this investigation to encourage exploration of a diverse set of methodologies, and groups tended to cluster around common methods. The resulting metadata reflect the most employed methods during the timeframe of this study (Figure. 1). For instance: nearly half of participants analyzing samples by 16S reported using the same DNA extraction kit (there were ∼15 other pre-identified options, as well as ‘In-house’ and ‘other’ possibilities); and only 2 labs (∼4%) used non-Illumina sequencing platforms.

### Comparing Results Between Laboratories

To help assess the impact of methodological choices in the context of compositionally-sensitive MGS measurements, we focused here on ratios between Phyla (e.g., the Firmicutes:Bacteroidetes ratio: Figure 5) instead of the raw relative abundances [46]. By using this strategy to remove the compositional dependence of MGS results, common statistical tools (e.g., mean, standard deviation, confidence intervals) could be directly applied. However, it must be noted that the Firmicutes:Bacteroidetes ratio only reveals the impact of particular methodological choices on the tested phyla (Firmicutes and Bacteroidetes). Thus, a methodological choice that only impacted Proteobacteria, as well as one that affected Firmicutes and Bacteroidetes similarly, would not be noted herein. Nevertheless, significant variability in the Firmicutes:Bacteroidetes ratio was observed (Figure 5) both between samples (presumably due to real differences between the samples) and between participating laboratories (presumably due to differences in measurement methodology).

When comparing between methodologies, the most basic experimental choice is between 16S and WGS, and this choice had further implications for how subsequent steps were performed (e.g., PCR conditions, library prep, sequencing depth, bioinformatic analysis). Thus, we first compared the Firmicutes:Bacteroidetes ratio between analysis methods (Figure 5). In this case, it turned out that the most basic choice of how to analyze samples had a statistically significant effect (Figure 6). Analysis of each of the stool samples individually (Figure SI-3) or statistical analysis using alpha diversity instead of the Firmicutes:Bacteroidetes ratio (Figure SI-4) confirmed the significant impact of analysis strategy on observed results. Practically, this raises real concerns about the comparability of data results between laboratories whose analyses differ between 16S and WGS analysis. More generally, researchers should use utmost caution when trying to compare between data sets collected using divergent experimental methods.

Within the data collected for the MSC, the significant effect observed for the choice of analysis strategy had the specific implication of further limiting statistical power (e.g., of the 44 participating labs, 30 reported 16S results and 14 reported WGS results). Nevertheless, the observed effects of other methodological choices could be similarly assessed for 16S (Figure 7) or WGS (Figure SI-4) results. Interestingly, while the statistical power was limited in this study, some methodologies still appeared to have either large effect sizes or large impacts on variability/precision. While it is tempting to draw firm conclusions from the current investigation, caution is warranted due to the limited sample sizes. Instead, it is hoped that this investigation will help guide further investigations.

### ‘Spike-in’ organisms

During production of the stool samples, two exogeneous, ‘spike-in’, whole cell bacterial strains were included, *A. fischeri* and *L. xyli*. Both strains are typically absent in human stool. With the addition of 10^8^ cells/mL, each organism was expected to comprise approximately 1 % of the total stool relative abundance, providing sufficient signal for identification without significantly affecting the overall sample profile. Unfortunately, while this expectation proved accurate for *A. fischeri, L. xyli* was not identified in any the of 16S MGS results and was only observed at a very low relative abundance by WGS (Figure SI-7). This absence or low-level detection could be the result of a number of sources including lack of representation in the databases, bias in the DNA extraction of *L. xyli*, or the amount of *L. xyli* added to the samples. However, multiple coauthors were individually able to reliably detect *L. xyli* using alternate bioinformatic pipelines, so it is likely that its limited detection in this dataset reflects a shortcoming in the reference database used [data not shown and manuscript in preparation]. This explanation is also supported by the observation that for WGS analyses, the variability of the ratio of spike-in relative abundances between samples was somewhat improved among the labs with the deepest sequencing results (Figure SI-2). It is worth noting that all raw fastq data submitted through the Mosaic Standards Challenge has been archived and made publicly available for the exploration of alternate bioinformatic methods.

The inability to reliably detect *L. xyli* within the framework of this project impacts our ability to accurately and confidently use *A. fischeri* as well since observing a constant ratio between the two spike-in organisms is fundamental to trusting their utility (Figure SI-7). Nevertheless, key considerations were identified for future experimental design and implementation of internal, spike-in controls. First, the strain should normally be absent in the sample, but still identifiable by the analysis/database used. This can be tricky because databases often focus on the organisms commonly encountered in each type of sample, and because the users of bioinformatic pipelines may not have easy access to the underlying reference databases at the time of analysis. Second, spike-in abundance should be sufficiently high that it can withstand potential losses in the processing and still be identified, while not significantly compromising the fraction of sample reads allocated the organisms native to each sample. This is in turn complicated by the dependence of the observed relative abundance of any spike-in organism on the MGS methods to be employed and their potential for bias with respect to each spike-in organism. And third, the inclusion of additional spike-in organisms (e.g., 3-4 spike-ins total) should be considered when MGS workflows have not been identified and tested a priori. This provides redundancy to accommodate wide ranges of MGS methodologies and biases. In this study, the inclusion of additional organisms could have avoided the problematic absence of *L. xyli* in the reference database.

### DNA Mock Communities

The DNA mixtures provided the ground-truth component in this study. Here, measurement bias was observed as a disagreement between the actual ratios (black bars show the 99 % confidence interval) and observed ratios (red and blue points) in Figure 8. This bias depends on both the particular taxa analyzed, as well as the methods employed (16S vs. WGS is broken out here). Interestingly, even where there was consensus between participating labs (i.e., a narrow boxplot indicating strong consensus), substantial bias was still observed (low accuracy). The consensus between participating labs is particularly apparent in the WGS analysis of the equi-genomic DNA mixtures (upper right panel, Figure 8) suggesting some systematic bias affecting each lab. Of note, these mixtures were comprised of genomic DNA from a prototype reference material. Since the time of the MSC, NIST has completed a full characterization of DNA from 19 bacterial strains; NIST Reference Material, RM 8376, is now available for researchers to construct their own DNA based mock communities [49].

## Conclusion

From 2017 to 2020, the MSC provided a set of biologically-derived and mock community microbiome samples, at no charge, to any interested MGS research group in an effort to identify the extent of methodological variability between researchers and assess its impact on measured taxonomic profiles. 44 research groups submitted both raw MGS data and detailed metadata about their in-house sample-handling protocols; although this represents a large number of participating laboratories by most interlaboratory efforts, it remained statistically limiting for the large number of metadata parameters (≈100) that were explored. Initial choices about analysis strategy (i.e., amplicon vs. shotgun) significantly impacted the observed Firmicutes:Bacteroidetes ratios across all samples. The null hypothesis of no significant effect could not be ruled out for most methodological choices within this study, though some appeared to have real effects on results (i.e., bias) or measurement precision (i.e., variability). Thus, the results collated herein should help refine the scope of future assessments of methodological choices. To this end, researchers at NIST have undertaken a pairwise approach to systematically compare select steps within the metagenomic workflow (manuscript in preparation). Additionally, through the inclusion of DNA mock communities with independently-measured ground-truth abundances, we were able to assess the accuracy of MGS measurements and observe significant and systematic measurement bias, even when participating laboratories achieved similar results. Overall, the MSC effort has significantly expanded our understanding of the impact of methodological choices on MGS measurement results and precision.

## Methods^1^

### Selection of Stool Donors

A total of 5 donors were selected from a donor pool maintained by TBC. Figure 2 shows a Bray-Curtis PCoA ordination plot the entire donor pool, including the 5 donors selected, based on their gut microbiome composition. The 5 donors were selected based on the dissimilarities of their microbiome composition (Figure 2).

### Sample Collection and Processing

All stool samples were collected in accordance with TBC’s Institutional Review Board protocol and have been de-identified. The donors were provided with collection kits, and samples were returned to the TBC via overnight shipping for processing. Upon receipt, the samples were aseptically transferred to a zip-top bag for dispensing. The samples were stored at -80 °C in 30 g aliquots until further processing. Multiple bowel movements were collected and pooled from each donor. Material from each donor was processed individually (to avoid cross contamination) and inside a biological safety cabinet. Using a Ninja blender, 150 g of fecal material was combined with 150 g to 300 g of dry ice and homogenized into a fine powder. The blender was loosely covered with a sterile lab tissue and placed in a -20 °C freezer overnight to allow the remaining dry ice to sublime. For each sample, before the addition of OMNIgene Stabilizing Solution (OGS), 50 g of neat powder was set aside and stored at -80 °C. Approximately 90 g of stool powder was added to 750 mL of OGS. The solution was covered and left to stir overnight at room temperature. The following morning, 1 mL aliquots were prepared and stored at –80 °C.

### Addition of Spike-In Bacteria and Aliquoting of Samples

Spike-in bacteria, *Aliivibrio fischeri* (formerly known as *Vibrio fischeri*, Gram negative) and *Leifsonia xyli* (Gram positive), were grown to an approximate density of 10^8^ CFU/mL and 10^9^ CFU/mL, respectively. Cell concentration was confirmed via plate count and optical density. The spike-in bacteria were concentrated by centrifugation, resuspended, and added to each stool solution 1 hour prior to aliquoting to ensure thorough homogenization. Working in a biological safety cabinet, the solution was aliquoted using wide-bore pipette tips into (800 to 850) aliquots. Final concentration of stool after addition of the spike-in was 100 mg/mL and final concentration of each spike-in organism was 10^8^ CFU/mL. Samples were stored at -80 °C until distribution.

### Sample QC

To assess the homogeneity of the stool samples, ten aliquots from each donor pool were subjected to 16S rRNA amplicon sequencing and shotgun metagenomic sequencing. All sample processing, DNA extraction, library preparation and sequencing steps were conducted at CosmosID (Germantown, MD) using proprietary protocols. For the 16S sequence data, reads were demultiplexed using split_libraries.py with default filtering parameters. 16S rRNA gene sequences were then sorted based on sample ID using the QIIME script *extract_seqs_by_sample_id*.*py*. Bacterial operational taxonomic units were selected using *pick_open_reference_otus*.*py* workflow. 16S rRNA taxonomy was defined by ≥ 97 % similarity to reference sequences using the *core_diversity_analyses*.*py script*. Alpha diversity, alpha rarefaction curves, and taxonomy assignments were determined using the core_diversity.py workflow. Data were rarefied to 100,000 sequences per sample to minimize the effect of disparate sequence number on the results. Alpha diversity metrics were computed from the average of 100 iterations from the alpha collated results. Microbiome features were quantified from metagenome data using existing [Metaphlan2, HUMAnN2, etc.] and in-house pipelines to identify strain-level taxonomic markers for all samples.

### DNA Mixtures

Mixtures of purified genomic DNA from thirteen ATCC-derived strains were prepared in 1X TE buffer at a final concentration of ≈100 ng/μL. The two mixtures were made by combining the genomic DNA from the following bacterial strains: *Staphylococcus aureus* ATCC BAA 44, *Staphylococcus aureus* ATCC 12600, *Pseudomonas aeruginosa* ATCC BAA 47, *Enterococcus faecalis* ATCC 19433, *Salmonella enterica* ATCC 700720, *Salmonella enterica* ATCC 12324, *Escherichia coli* ATCC 43895, *Staphylococcus epidermidis* ATCC 12228, *Klebsiella pneumoniae* ATCC 13883, *Shigella sonnei* ATCC 25931, *Streptococcus pyogenes* ATCC 12344, *Corynebacterium amycolatum* ATCC 49386. The individual genomic DNA components were part of a prototype reference material and were not fully characterized at the time of the MSC. Subsequent analysis revealed some of the materials were cross contaminated with other components from the prototype materials including *Achromobacter xylosoxidans*. Mix A was designed to be equi-genomic with calculated relative abundances by mass of each strain ranging from ≈6.8 % to ≈10 %. Mix B was designed as a log-dilution of the genomes varying across 3 orders of magnitude (from ≈0.01 % to ≈30 % by mass). For each mixture (A and B), we prepared a single pool and then distributed across 700 aliquots where each contained approximately 20 μL (2 μg) of DNA per aliquot. An average (across all participating laboratories) relative abundance plot for each sample by amplicon or shotgun sequencing is included in the supporting information (Figure SI-8). We performed digital droplet PCR (ddPCR) to measure the absolute abundance as ground truth for the following species in the mixture: *Enterococcus faecalis, Klebsiella pneumoniae, Pseudomonas aeruginosa*, and *Streptococcus pyogenes*. These species were selected because they were taxonomically distinct within the mixtures at the Genus level, facilitating MGS discrimination. Pairwise ratios of these abundances provided the ground truth values depicted in Figure 8. The validated ddPCR assays were reported previously [49].

### Interlaboratory Study Executio

#### Recruitment

Starting in the Spring of 2018, we launched a media campaign that targeted the scientific community via social media and email blasts as an attempt to recruit a large and international cohort of participants. After the MSC launched in May 2018, we continued the outreach campaign via public speaking engagements at various international microbiology conferences. We actively recruited volunteers up until January 2020 when the MSC officially closed. MSC reference materials were shipped to any lab in the world, upon request, from May 2018 till January 2020.

#### Data Availability

The Mosaicbiome.com web portal was used during MSC to store, analyze, visualize, and share all the raw data and metadata that was submitted by the MSC participants. However, in the Spring of 2022, the site was discontinued due to the costs associated with data storage and maintenance. Therefore, these data (fastq files and metadata summaries) have been made available via https://data.nist.gov. [exact link TBD]

#### Sample Availability

At the time of publication, many aliquots of the stool and DNA materials generated through the Mosaic Standards Challenge still remain available from TBC.

### Taxonomic Profiling of Interlab Data

All raw sequence data (fastq files) generated by interlab participants were downloaded from the MosaicBiome web portal and subsequently analyzed via the CosmosID (www.CosmosID.com) taxonomic classification tool using the CosmosID reference genome databases (WGS version: 1.0.2; 16S version: 1.1.0). The MGS results (taxonomic profiles) for all the MSC data are publicly available and can be found by visiting https://app.cosmosid.com and following the directory structure: *Datasets -> Example_Datasets -> Mosaic_Microbiome*.

### Analysis of mosaic data results and methodological parameters

A total of 50 datasets were received. Two datasets were dropped due to incomplete metadata and an additional 4 labs were The MGS results (taxonomic profiles) and associated metadata from all submitted data sets were analyzed using R. The raw data and code used for analysis and to generate the figures in this manuscript have been shared via https://data.nist.gov. [exact link TBD]

## Supporting information

Metadata Worksheet

## Acknowledgements

Funding for the production of the fecal reference materials and reference material shipping was generously provided by the Janssen Human Microbiome Institute (JMHI).

## Footnotes

1 Certain commercial equipment, instruments, or materials are identified in this paper to foster understanding. Such identification does not imply recommendation or endorsement by the National Institute of Standards and Technology, nor does it imply that the materials or equipment identified are necessarily the best available for the purpose. The reference materials used in this study were not certified by NIST and are not official <u>NIST Reference Materials</u>.

