## Supplementary material for "Variability and Bias in Microbiome Metagenomic Sequencing: an Interlaboratory Study Comparing Experimental Protocols": SI Figures

### External Mosaic SI Figures

- **Metadata reporting document**: This spreadsheet includes the metadata that was requested from all labs submitting FASTQ results. Many questions had pre-populated answer options to select from, while other quesitons had text-input fields. ([link]())

| Main category | Subcategory | Child question? | Amplicon | WGS | Preferred |
|---|---|---|---|---|---|
| Sequencing type | Sequencing type | SUPER-PARENT | x | x | x |
| Sample storage | Sample_storage | PARENT | x | x | x |
| | Sample_storage | CHILD | x | x | x |
| | Sample_storage | CHILD | x | x | x |
| | Sample_storage | CHILD | x | x | |
| | Sample_storage | | x | x | |
| DNA extraction | DNA_extraction | | x | x | x |
| | DNA_extraction | | x | x | x |
| | DNA_extraction | | x | x | |

- **Figure SI-1**: Metagenomic sequencing analysis of Mosaic fecal samples to determine homogeneity of samples. The bar chart shows the relative abundance as measured by WGS MGS at the genus level for 10 replicate tubes from each stool sample (Stool 1-5). Taxa colors denote the 15 most abundant genera overall, as well as the two exogeneously taxa; all other genera are grouped as 'other' and shown in grey. MGS analysis by 16S is shown in Figure 3 of the main manuscript.

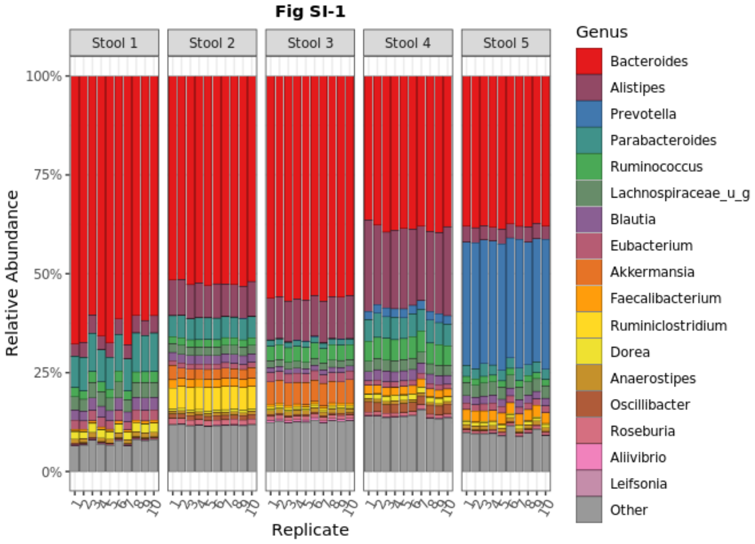


- ○

- **<u>Figure SI-2</u>**: The read depth varied substantially between unique lab submissions and even among samples analyzed within a lab. The total counts across all taxa are plotted here for each lab and sample reporting 16S amplicon sequencing (A) or shotgun metagenomic sequencing (B) results.

**Figure SI-2A**

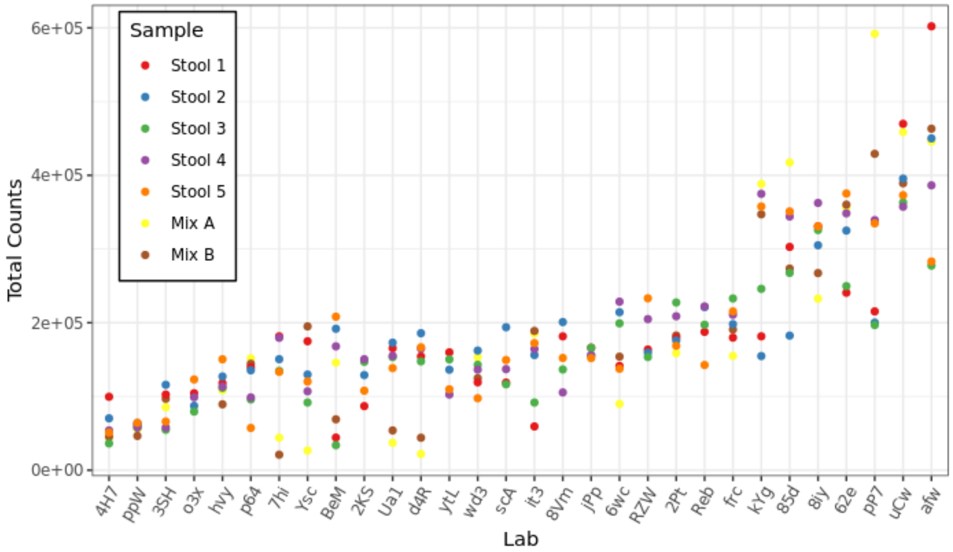


  - 

**Figure SI-2B**

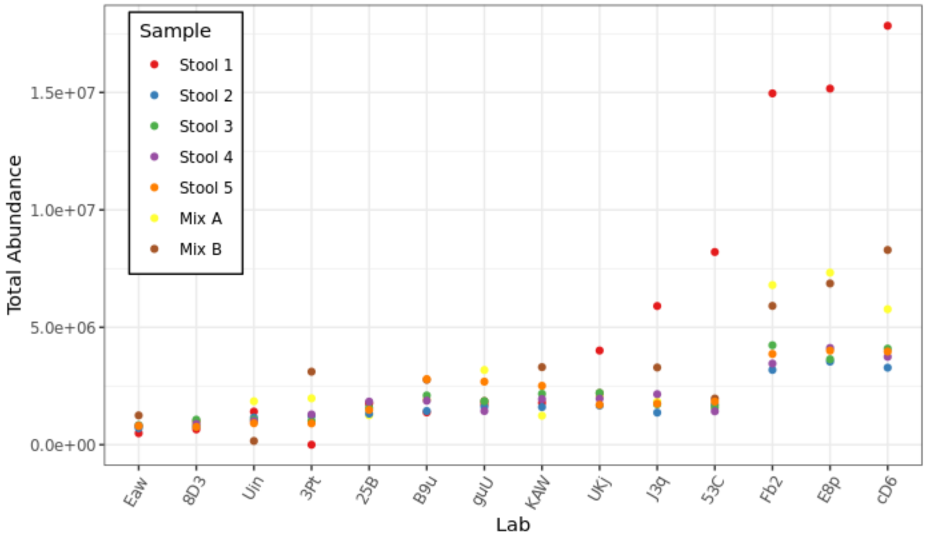


  - 

- **<u>Figure SI-3</u>**: The effect analysis strategy (16S versus WGS) on the Firmicutes:Bacteroidetes ratio was quantified for each Stool sample by dividing the average results among labs reporting the specified parameter level by the average results overall. For each sample, this parameter effect was plotted on a log (base 2) scale, such that a horizontal line at 0 denotes the null hypothesis of no effect; error bars show the 99% confidence interval. Analysis of a single stool sample is included in the main manuscript (Figure 5), and analyses the other four stool samples are included here (Figures SI-3A, SI-3B, SI-3C, and SI-3D) and show similar results. Similar stratification was also observed when measuring each sample's Inverse Simpson alpha diversity (Figure SI-4).

○ 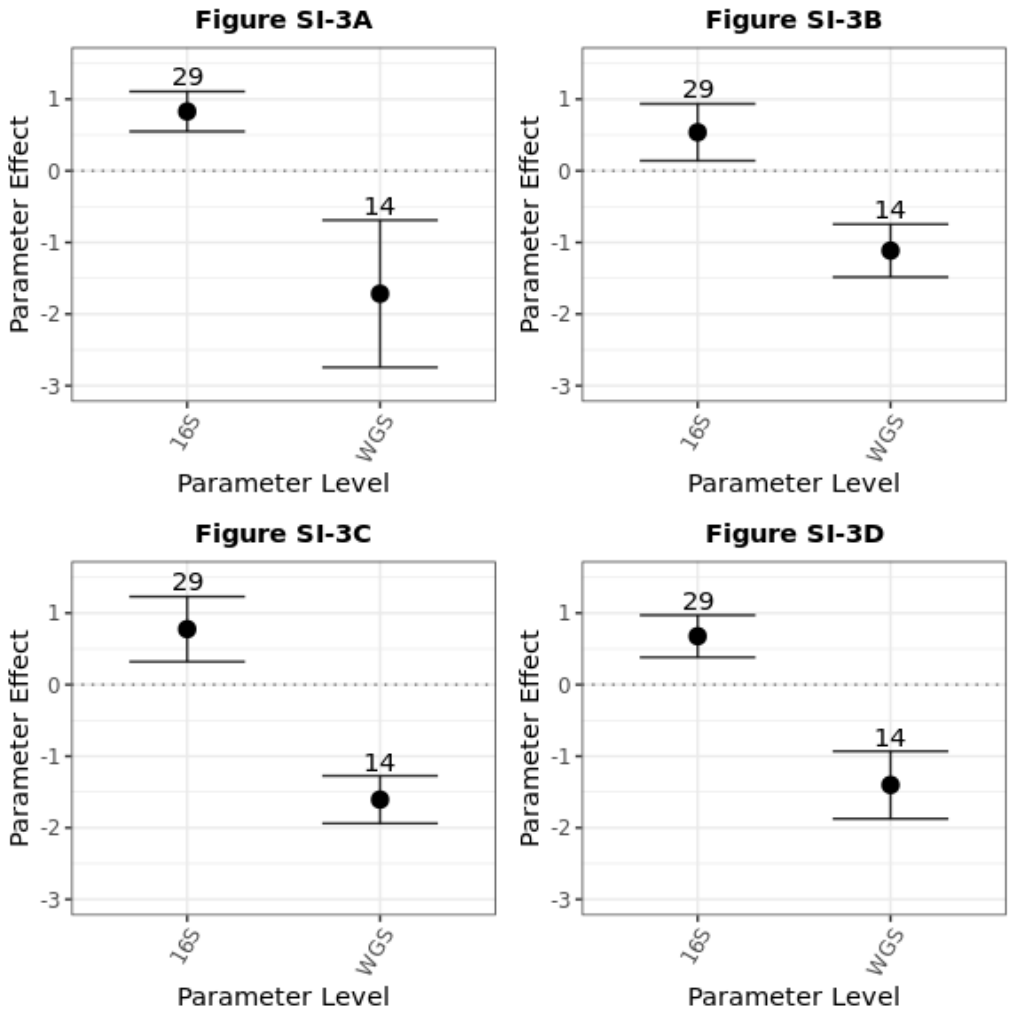


○

- **<u>Figure SI-4</u>**: The effect analysis strategy (16S versus WGS) on alpha diversity was quantified. For each sample, the parameter effect was calculated by dividing the average Inverse Simpson alpha diversity among labs reporting each specified parameter level by the average results overall. The parameter effects were plotted on a log (base 2) scale, such that the horizontal lines at 0 denote the null hypothesis of no effect; error bars show the 99% confidence interval. Analyses of all five stool samples (Figures SI-4A, SI-4B, SI-4C, SI-4D, SI-4E) exhibit similar stratification with analysis strategy as analyses of the Firmicutes:Bacteroidetes ratio (as shown in Figure 5 of the main manuscript and Figure SI-3).

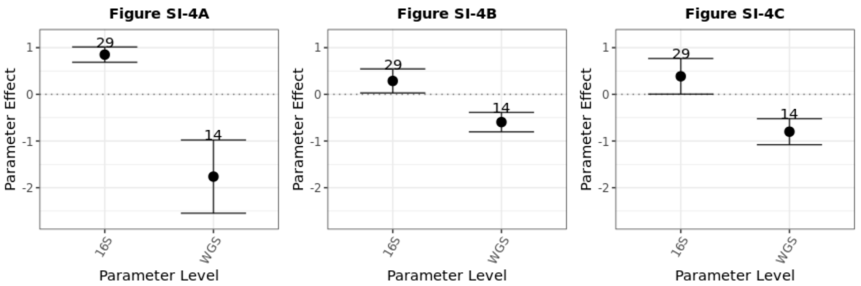


○

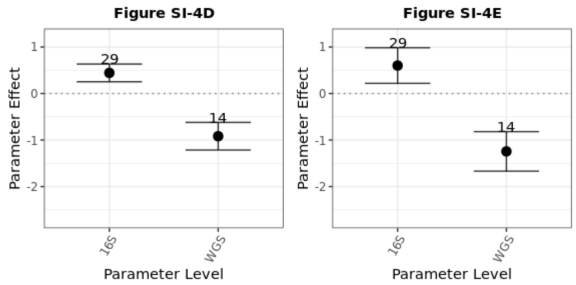


- **<u>Figure SI-5</u>**: Within labs performing 16S amplicon sequencing, the parameter effect on the Firmicutes:Bacteroidetes ratio was calculated as described in Figure 5 for each relevant metadata parameter. Shown in Figure 6 for just one stool sample, results from the other 4 stool samples are included here (Figure SI-5A, SI-5B, SI-5C, SI-5D). Parameter levels here are identical to those defined in Figure 6.

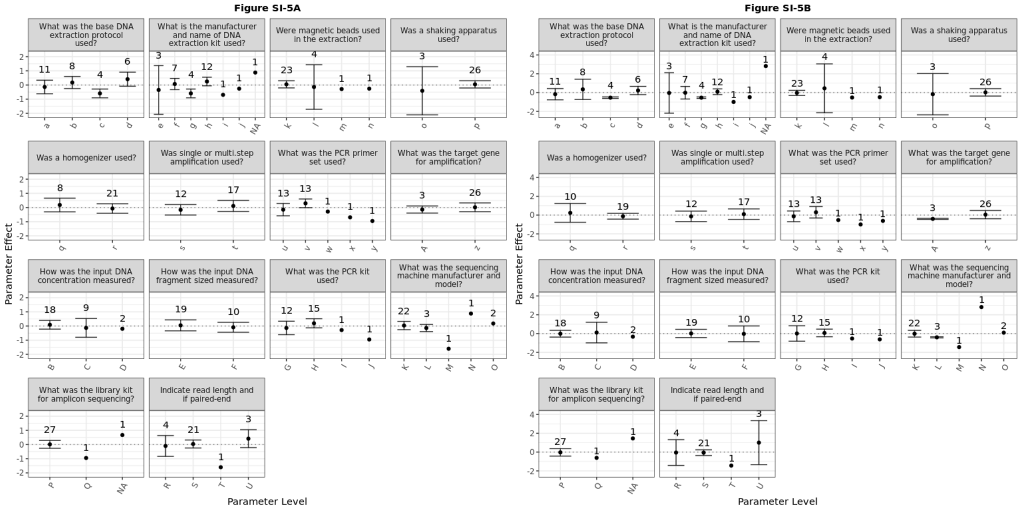


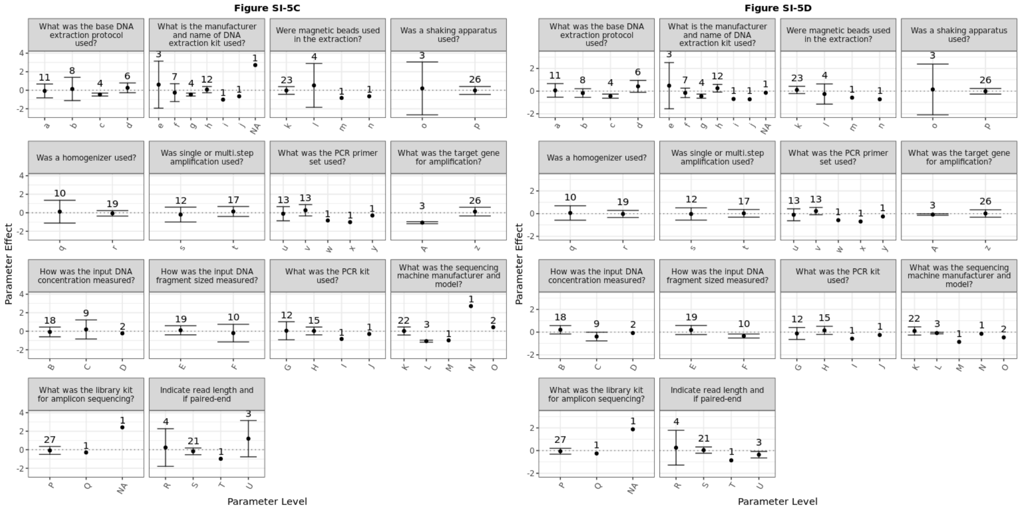


- **<u>Figure SI-6</u>**: Within labs performing WGS amplicon sequencing, the parameter effect on the Firmicutes:Bacteroidetes ratio was calculated as described in Figure 5 for each relevant metadata parameter and each of the five stool samples (Figure SI-6A, SI-6B, SI-6C, SI-6D, SI-6E). Parameter levels here are defined in the associated table.

Figure SI-6A

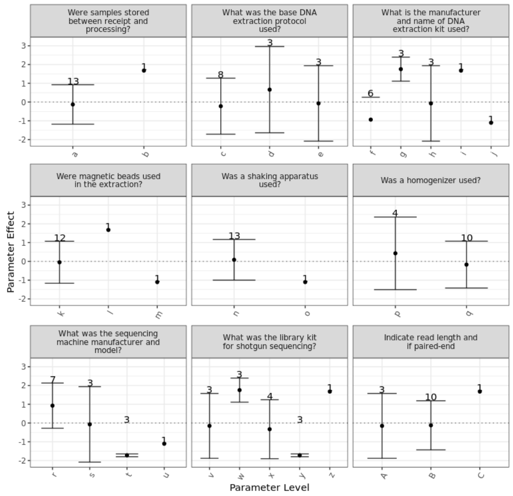


Figure SI-6B

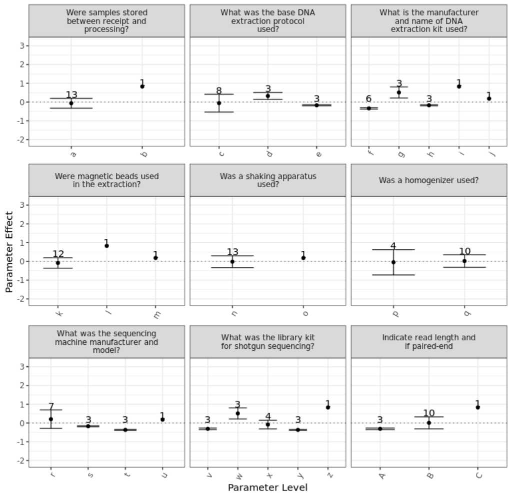


○

Figure SI-6C

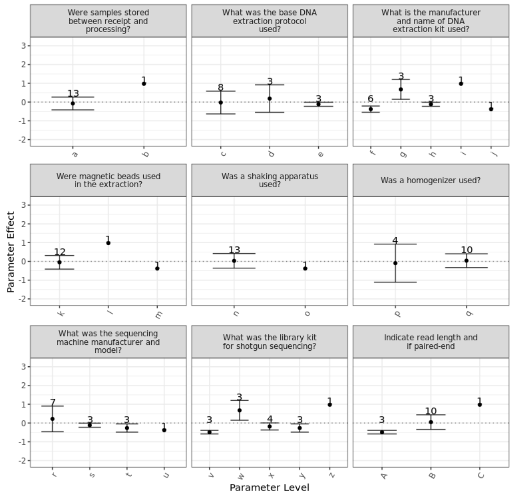


Figure SI-6D

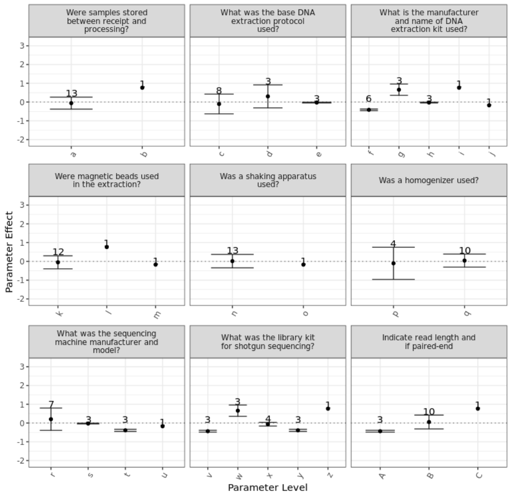


Figure SI-6E

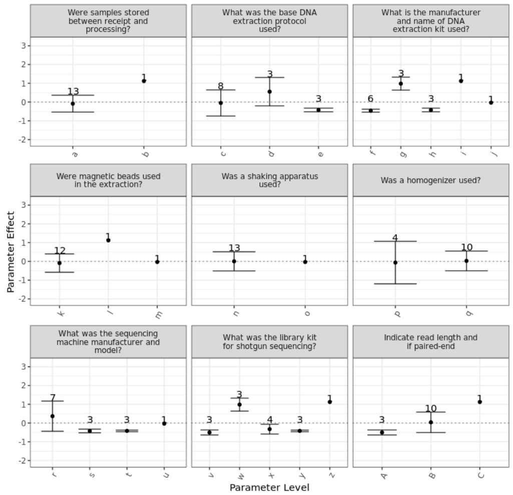


| Symbols | Parameter | Option |
|---|---|---|
| a, b | Were samples stored between receipt and processing? | (a) Yes, (b) No |
| c, d, e | What was the base DNA extraction protocol used? | (c) Others, (d) Internal, (e) HMP |
| f, g, h, i, j | What is the manufacturer and name of DNA extraction kit used? | (f) No-kit Other method, (g) PowerMag, (h) Mo Bio DNeasy PowerSoil Kit, (i) Zymo fecal DNA Miniprep kit; ZR96 fecal DNA kit, (j) In-house protocol |
| k, l, m | Were magnetic beads used in the extraction? | (k) glass_beads_0,1mm, (l) No, (m) zirconia_beads_0,1mm |
| n, o | Was a shaking apparatus used? | (n) Yes, (o) No |
| p, q | Was a homogenizer used? | (p) No, (q) Yes |
| r, s, t, u | What was the sequencing machine manufacturer and model? | (r) Illumina NextSeq 500, (s) Illumina HiSeq 4000, (t) Illumina MiniSeq, (u) Illumina NovaSeq |
| v, w, x, y, z | What was the library kit for shotgun sequencing? | (v) Illumina Nextera DNA, (w) Other, (x) Illumina Nextera XT, (y) NEB NEBNext Ultra DNA, (z) Kapa DNA Library |
| A, B, C | Indicate read length and if paired-end | (A) 150 PE, (B) 250 PE, (C) 300 PE |

- **Figure SI-7**: Whole cells from two exogeneous taxa (*A. fischerii* and *L. xyli*) were added uniformly to each stool sample. The ratio of their observed relative abundances in each sample (by WGS MGS) is plotted here. While these ratios were expected to vary between labs due biases inherent to the different methods employed, high reproducibility was expected between samples within a single lab due to the constant actual abundances in each stool sample. Instead, substantial variability was observed both between labs and between samples.

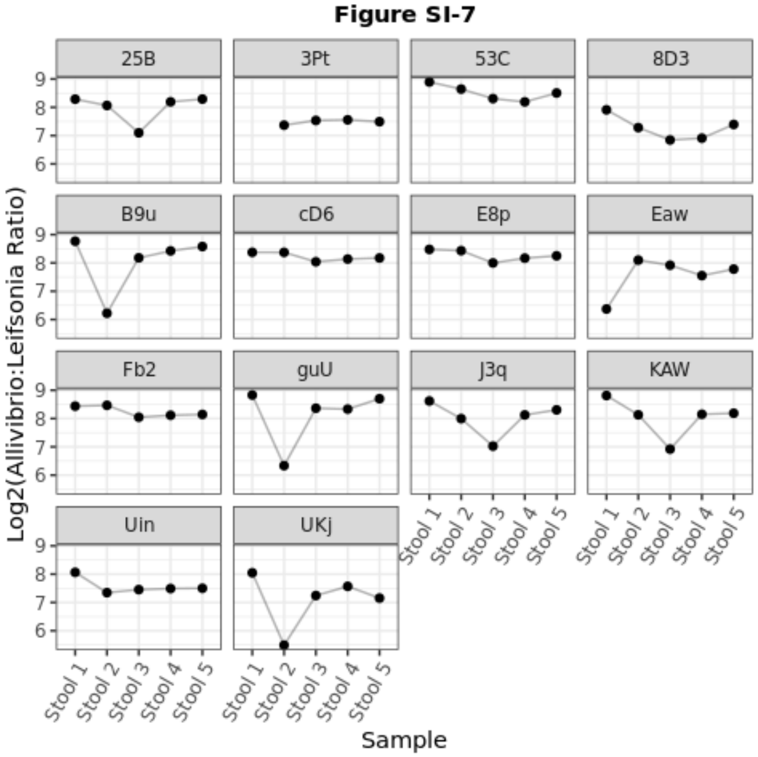


○

- **Figure SI-8**: Genus-level taxonomic bar charts depict the average observed composition (across all participating laboratories) of the DNA mock communities as observed through 16S (A) and WGS (B) MGS analyses. Taxa colors denote the most abundant Genera detected overall; taxa at relative abundance ≤ 0.4% are grouped as 'other' and shown in gray).

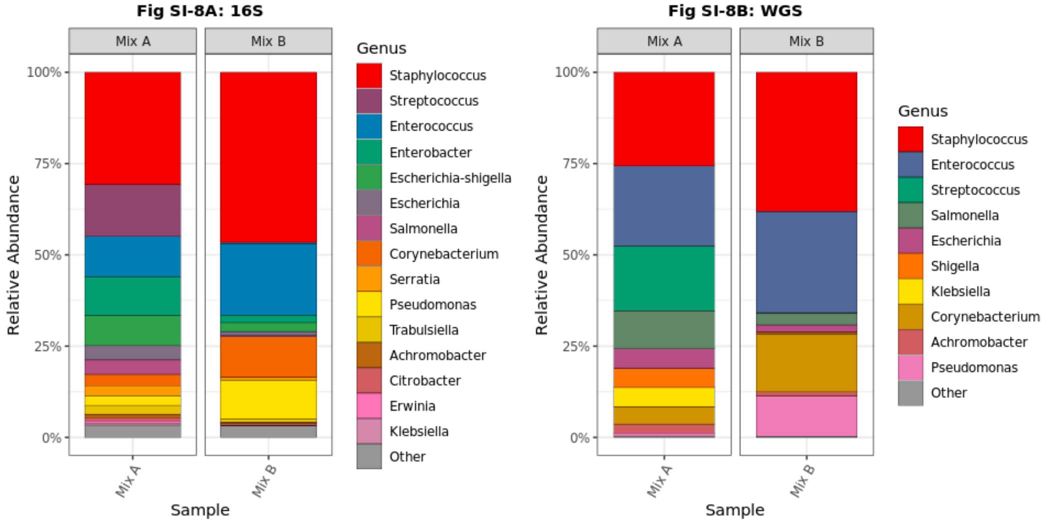


○
